# Gene tree discordance generates patterns of diminishing convergence over time

**DOI:** 10.1101/059006

**Authors:** Fabio K. Mendes, Yoonsoo Hahn, Matthew W. Hahn

## Abstract

Phenotypic convergence is an exciting outcome of adaptive evolution, occurring when species find similar solutions to the same problem. Unraveling the molecular basis of convergence provides a way to link genotype to adaptive phenotypes, but can also shed light on the extent to which evolution is repeatable and predictable. Many recent genome-wide studies have uncovered a striking pattern of diminishing convergence over time, ascribing this pattern to the presence of intramolecular epistatic interactions. Here, we consider gene tree discordance as an alternative driver of convergence levels over time. We demonstrate that gene tree discordance can produce patterns of diminishing convergence by itself, and that controlling for discordance as a cause of apparent convergence makes the pattern disappear. We also show that synonymous substitutions, where neither selection nor epistasis should be prevalent, have the same diminishing pattern of molecular convergence among closely related primate species. Finally, we demonstrate that even in situations where biological discordance is not possible, errors in species tree inference can drive these same patterns. Though intramolecular epistasis is undoubtedly affecting many proteins, our results suggest an additional explanation for this widespread pattern. These results contribute to a growing appreciation not just of the presence of gene tree discordance, but of the unpredictable effects this discordance can have on analyses of molecular evolution.

## 1. Introduction

Understanding the molecular basis of adaptation is a central goal in evolutionary biology. Among the many strategies used to address this problem, the study of convergence – when different species solve similar evolutionary problems using similar solutions – provides a unique look into adaptive evolution. More specifically, comparing the molecular causes of convergence among taxa can reveal how repeatable and predictable adaptation is, and concomitantly clarify the link between adaptation and genotype (Rosenblum et al. 2014; Storz 2016)

Motivated by these goals, recent studies have been successful in demonstrating that convergent molecular changes often accompany convergent phenotypes (e.g., Christin et al. 2008; Manceau et al. 2010; Dobler et al. 2012; Feldman et al. 2012; Zhen et al. 2012; Projecto-Garcia et al. 2013), a fact that historically was not always accepted (Stern 2013). However, these studies are still not informative as to how frequently molecular convergence is truly adaptive, nor of how widespread it is in nature. Fortunately, the availability of whole genomes has allowed researchers to also examine the frequency of molecular convergence across all proteins. These comparative studies have reported high levels of molecular convergence in many clades (e.g., Bazykin et al. 2007; Rokas and Carroll 2008; Parker et al. 2013; Foote et al. 2015; Goldstein et al. 2015; Zou and Zhang 2015a), although discerning the evolutionary (neutral vs. adaptive) and molecular mechanisms driving these substitutions is still difficult. Among many reasons for this difficulty, the uncertainty in the choice of a null model in studies of adaptive convergence has cast doubt on multiple previous results (Thomas and Hahn 2015; Zou and Zhang 2015b).

Inferring the specific molecular mechanisms underlying convergence is difficult, and may be intractable from genome data alone. However, a number of studies have used patterns of molecular convergence to infer general molecular mechanisms underlying protein evolution. One pattern common to many studies of convergence is the reduction in molecular convergence between species over time (e.g., Rogozin et al. 2008; Povolotskaya and Kondrashov 2010; Naumenko et al. 2012; Goldstein et al. 2015; Zou and Zhang 2015a). This pattern has been suggested to be due to epistasis: substitutions at one site in a protein constrain the set of allowable amino acids at other sites (Zou and Zhang 2015a) As species diverge, amino acid substitutions that were once forbidden become allowed (and vice versa), reducing the probability that similar evolutionary problems will be solved by the same amino acid substitution in multiple lineages (Goldstein et al. 2015). Over time the frequency of convergence would therefore decrease, and similar arguments have been made about the frequency of reversion over time (e.g., McCandlish et al. 2016).

Previous studies of molecular convergence (adaptive or not) have understood these patterns to represent molecular homoplasy; that is, substitutions are assumed to have happened independently along the branches leading to the two or more taxa that converged to the same character state. There is, however, a largely overlooked source of apparent molecular convergence that does not match these assumptions: gene tree discordance. Discordance causes the false appearance of convergence when no independent substitutions have actually occurred (Hahn and Nakhleh 2016; Mendes and Hahn 2016; Storz 2016).

Gene tree discordance, also known as phylogenetic incongruence, occurs when phylogenies obtained from individual genes differ among themselves and from the species phylogeny. This phenomenon can be caused by diverse biological factors such as incomplete lineage sorting (ILS), introgression, and lateral gene transfer (Maddison 1997). With the increasing availability of sequences from whole genomes and across clades, it has become evident that gene tree discordance is common in nature, due to both ILS (e.g., Pollard et al. 2006; Scally et al. 2012; Brawand et al. 2014; Jarvis et al. 2014; Zhang et al. 2014; Suh et al. 2015; Pease et al. 2016) and introgression (reviewed in Mallet et al. 2016). The extent of ILS (and consequently of gene tree discordance) is positively dependent on ancestral population sizes and inversely dependent on the time between speciation events (Pamilo and Nei 1988; Maddison 1997), but is independent of timescale, affecting nodes regardless of how old they are (Oliver 2013).

Phylogenetic incongruence can produce apparent convergence by means of hemiplasy, a term coined to signify the production of a homoplasy-like pattern by a non-homoplasious event (Avise and Robinson 2008). At the sequence level, hemiplasy has been recently dubbed “SPILS” (Substitutions Produced by ILS) because of the repeatable and predictable pattern of substitutions it produces under ILS (Mendes and Hahn 2016). SPILS is expected to occur whenever substitutions along discordant internal branches (i.e., internal branches on discordant gene trees that are absent in the species tree) are resolved on the species tree (Fig. 1). SPILS will artificially create molecular convergence (or reversions) because substitutions along discordant internal branches will appear to have occurred multiple times when these substitutions are mapped to the species tree (e.g., site 1 in Fig. 1). Hemiplasious molecular convergence does not reflect any biological property of the sites where it occurs, being solely the result of the technical bias resulting from using a species tree that does not match the topology of the gene or region being analyzed.

**Figure 1.**
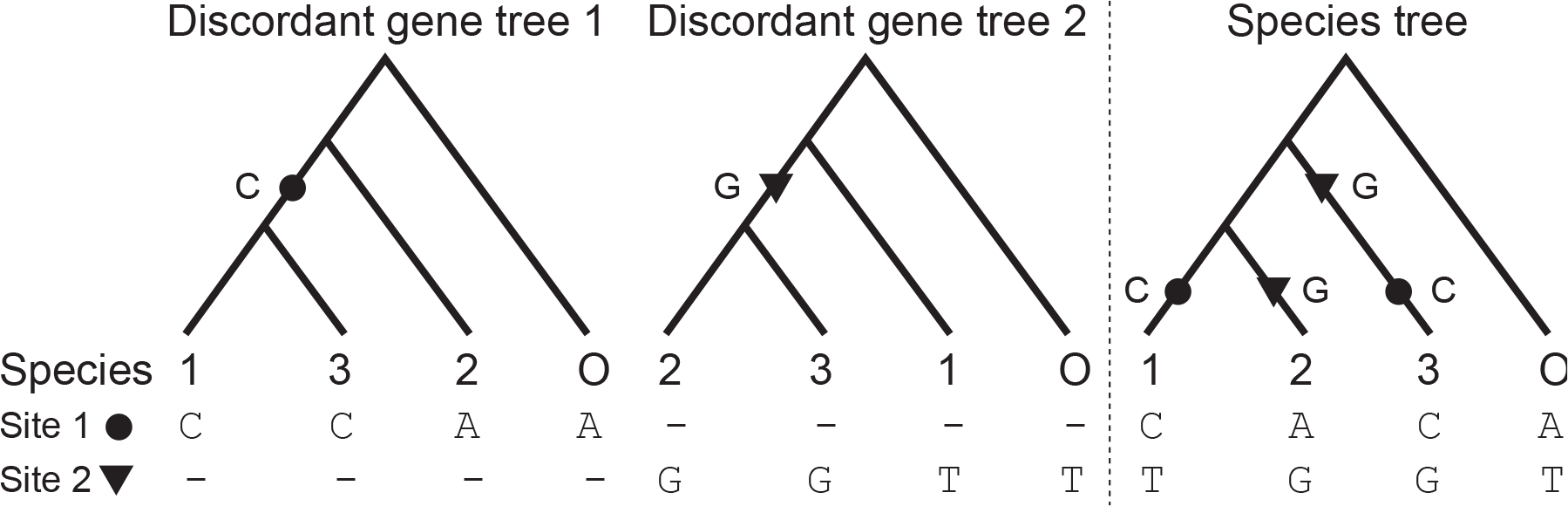
Two discordant gene trees and their species tree. The circle and triangle on discordant gene trees 1 and 2 correspond to substitutions at two different, loci., from an ancestral allele to C and, G., respectively. If either gene is analyzed under the species, tree., their substitutions (circle or triangle) are incorrectly mapped on leaves (species 1 and 3, or 2 and 3) rather than on internal, branches., and interpreted as molecular convergence.

Because both ILS (Pamilo and Nei 1988) and introgression (Coyne and Orr 1989) are more likely between more closely related species, we predicted that discordance could produce declining patterns of molecular convergence similar to those observed previously. This source of apparently declining convergence would not be a consequence of epistasis, though of course epistatic interactions could be occurring in all proteins. One important distinction that can help us to distinguish between the causes of apparent molecular convergence is between “parallel” and “convergent” substitutions (Zhang and Kumar 1997) Substitutions are classified as parallel when the lineages being compared have the same ancestral state (Fig. 2a), and are classified as convergent if they have different ancestral states (Fig. 2b). Importantly, SPILS will only ever lead to the appearance of parallel substitutions, allowing us to separately consider a class of convergent substitutions that cannot be due to discordance alone.

**Figure 2.**
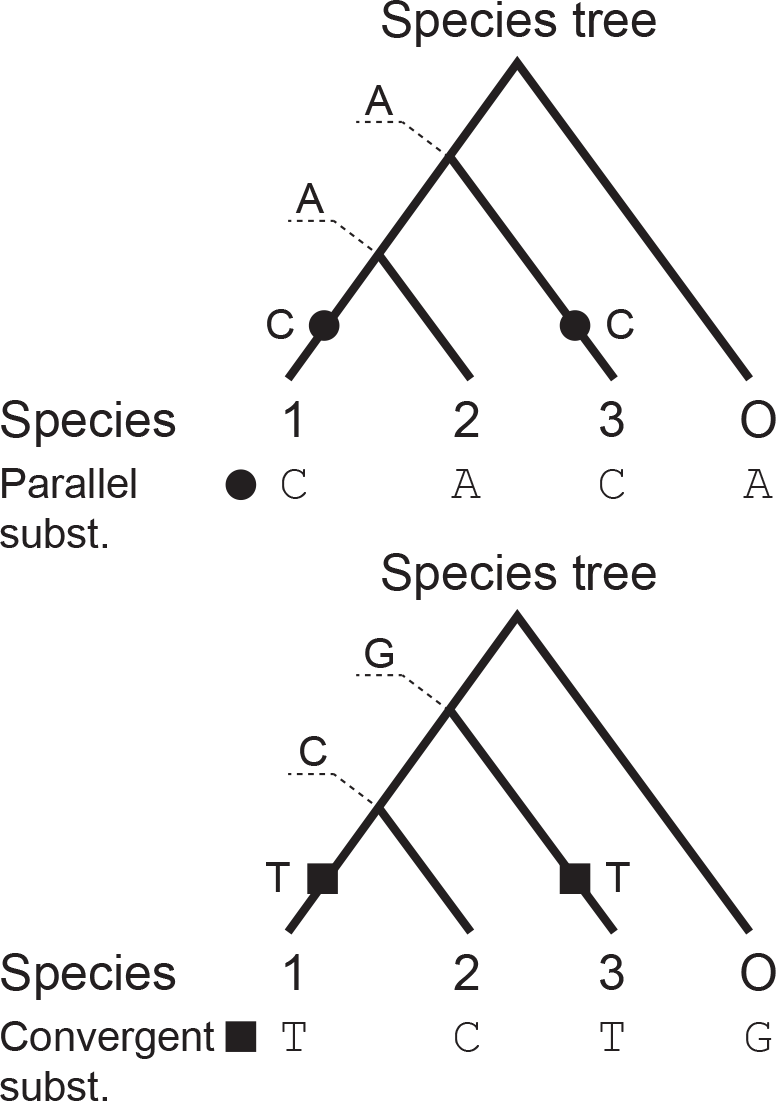
Sites undergoing molecular convergence can be classified as either parallel (top tree) if ancestral states are the, same., or convergent (bottom panel) if otherwise.

In what follows, we examine and control for the impact of varying levels of discordance on observed molecular convergence in multiple primate genomes, as well as in sequences simulated with ILS. Furthermore, we investigate whether substitutions that are unlikely to be constrained by epistasis (at synonymous sites) also exhibit decaying levels of convergence with time, presumably solely as a result of gene tree discordance. Finally, we examine errors in species tree inference as yet another source of discordance that can lead to apparent convergence, even when ILS and introgression are not expected to occur.

## Results

### When discordance is controlled for, the same amount convergence is observed regardless of phylogenetic distance

Using coding sequences from 12 primate species and reconstructing ancestral states of individual nucleotides using PAML (Yang 2007), we recovered the same previously described pattern of decaying convergence (normalized by divergence) as a function of genetic distance (Fig. 3a; Zou and Zhang 2015a). Although the original authors’ hypothesis was that this pattern is due to intramolecular epistasis, we propose that it is due to gene tree discordance with the single species tree that all substitutions are analyzed on. As laid out in the Introduction, the logic here is that more closely related species will tend to have shorter internal branches separating them, which will lead to greater levels of either ILS or introgression, and in turn increase the effect of SPILS as a source of artifactual molecular convergence. Note that convergence on sister lineages is not considered, so the internal branches separating taxa is the main determinant of their genetic distance.

**Figure 3.**
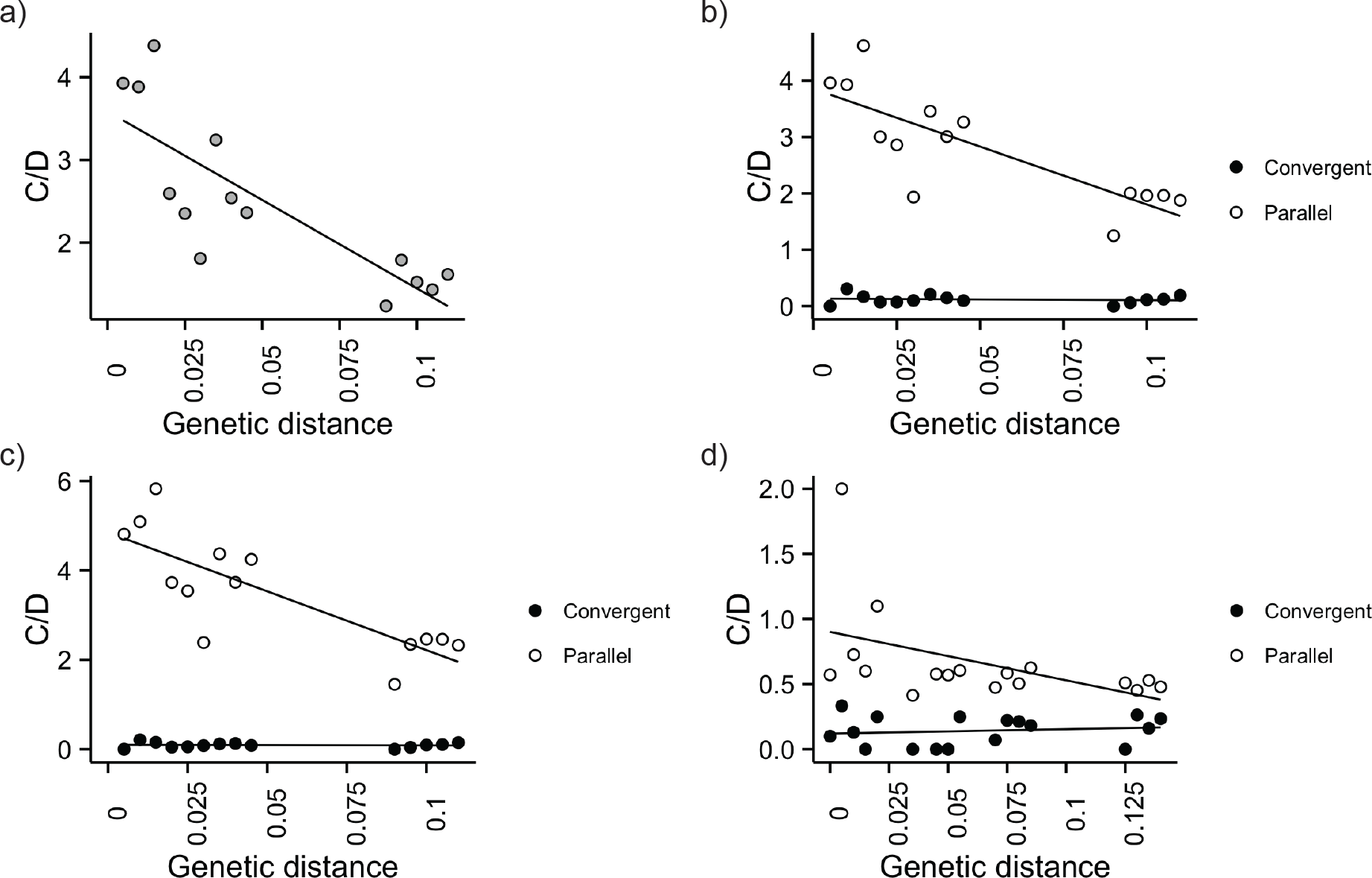
Convergence as a function of genetic distance for all pairs of primate species. a) All substitutions. b) Same as in panel a), but with parallel and convergent substitutions shown separately. c) Same as in panel a), but with only synonymous parallel and convergent substitutions shown. d) Sequences simulated on the primate species tree under the multispecies coalescent. In all panels the number of convergent substitutions (of whatever type) is normalized by the number of divergent substitutions (denoted C/D).

As discussed earlier, convergent substitutions produced by discordance will appear as “parallel” substitutions that have the same ancestral state, since there is truly only one substitution that has occurred (Fig. 2a). We can therefore control for discordance as a cause of the decreasing pattern of convergence by only considering “convergent” substitutions having different ancestral states (Fig. 2b). This is not to say that parallel sites will necessarily be caused by SPILS alone, only that by ignoring them we ensure that only true convergence is being analyzed.

Considering parallel and convergent sites separately in the primate data, we found that parallel sites exhibit a pattern of decaying convergence as a function of divergence, but that convergent sites do not (Fig. 3b). This implies that discordance is the major cause of the decreasing pattern of apparent convergence, and that epistatic interactions may play only a minor role in affecting the rate of truly convergent substitution. If discordance is controlled for, convergence between two species appears to be independent of the genetic distance separating them. Interestingly, we have found similar results reported for other datasets examining the decrease in convergence over time: only parallel substitutions decrease with time, while convergent substitutions show no relationship with time (see Figure 5 in Povolotskaya and Kondrashov 2010, and Figure 2a in Goldstein et al. 2015). Though neither of these studies discussed their results in light of discordance, we consider these analyses to lend further support to our hypothesis that SPILS is the true cause of these patterns. Alternatively, epistasis may affect parallel and convergent substitutions in a different manner. In order to address this hypothesis we next analyzed data where no epistatic interactions are expected.

### When epistasis is controlled for, convergence levels still decrease as a function of phylogenetic distance

We attempted to examine patterns of convergence in the absence of epistasis using two different approaches. We measured convergence levels (i) for synonymous substitutions within the same primate coding sequences analyzed above, as these sites should be less frequently involved in epistatic interactions; and (ii) in sequences generated through coalescent simulations that did not incorporate epistasis, but did allow for ILS (sequences were simulated along the same primate phylogeny; see Methods). If epistasis is the major reason why more closely related species exhibit greater levels of convergence, both of these approaches are expected to result in the same amount of convergence at all measured genetic distances.

Convergence levels at synonymous sites closely matched those measured when all sites were considered, with diminishing convergence as a function of time being observed only for parallel synonymous substitutions (Fig. 3c). While synonymous sites must occasionally be involved in epistatic interactions, we expect that the dataset containing only this site class harbors far fewer epistatic interactions, and should therefore exhibit convergence levels at least distinguishable from those found in the analysis of all nucleotide substitutions (which includes synonymous and nonsynonymous changes). The lack of such difference, however, suggests that the main driver of convergence levels is independent of the functional relevance of a site, and that it affects parallel substitutions, but not convergent substitutions. As explained above, the phenomenon of gene tree discordance as a result of ILS and introgression (coupled with hemiplasy) meets both criteria and nicely explains these results.

The same pattern of higher apparent convergence among more closely related species was observed in our simulated dataset, in which gene tree discordance occurs as a result of ILS, but where epistasis is absent (Fig. 3d). Furthermore, the decay of convergence was again observed only for parallel substitutions, as expected (Fig. 3d). These results provide further support for the role of gene tree discordance as a driver of convergence levels, as they suggest that processes that induce it (e.g., ILS) can be solely responsible for the higher apparent convergence that is observed among more closely related species. While this hypothesis is certainly most parsimonious in that it does not invoke epistasis, neither does it completely exclude a role for epistatic interactions.

### Tree inference error causes gene tree discordance and creates apparent convergence

Multiple studies have found the previously described reduction in convergence over time in cases where discordance among loci can clearly occur (e.g., Rogozin et al. 2008; Povolotskaya and Kondrashov 2010; Naumenko et al. 2012; Zou and Zhang 2015a). However, an additional study reported this difference using mtDNA genes and a species tree inferred from mtDNA (Goldstein et al. 2015). Such results are intriguing because although gene trees from the mitochondrial genome can disagree with the species tree obtained from nuclear genes, there should be no phylogenetic incongruence among mitochondrial genes nor between them and the mitochondrial species tree. The lack of recombination between mitochondrial loci should cause the mitochondrial genome to be inherited as a single topologically homogeneous genomic block. This result therefore appears to conflict with the hypothesis that discordance is the cause of these patterns, as apparent convergence due to SPILS should be impossible.

While gene tree discordance can be caused by ILS or introgression when multiple speciation events occur within a short time interval (i.e., when species tree internal branches are short), such circumstances should also be particularly permissive to phylogenetic incongruence as a result of tree inference error. The shorter an internal branch is, the less phylogenetic information it will contain, and so the greater are the chances of incorrectly inferring the relationship between taxa. It then follows that because more closely related lineages will tend to be separated by shorter internal branches, their placement will more likely differ due to errors in tree inference. As a result, even in datasets where biologically generated discordance is impossible, we may observe more apparent convergence because of SPILS (note that the term SPILS does not apply only when ILS is present, but whenever spurious substitutions are produced because of gene tree discordance; Mendes and Hahn 2016).

Importantly, Goldstein et al. (2015) inferred the species tree containing over 600 vertebrates using only a single mtDNA gene, slightly more than 500 amino acids long. They then analyzed the remaining mtDNA genes on this species tree. Although many relationships inferred among lineages were unresolvable using this dataset (i.e., had a branch length of zero), the species tree used is fully bifurcating. In effect, this means that the tree inference software has made arbitrary decisions regarding the relationships among some taxa, with the largest effect on closely related taxa.

In order to mimic the data used in Goldstein et al. (2015) and to test our predictions about the causes of discordance (and therefore convergence), we used the Jukes-Cantor model (Jukes and Cantor 1969) and the large species tree from this study to simulate 100 1-kb-long nucleotide sequences, again in the absence of epistasis. These simulations were done along a static species tree (i.e., they did not incorporate the coalescent process and were, as a result, “ILS-free”), and did not include within-gene recombination. Thus, all sites from all simulated loci had the same underlying genealogy – any discordance among inferred gene trees would have to be caused by errors in phylogenetic tree reconstruction. Note that here we use the term gene tree discordance to mean topological differences between gene trees, regardless of whether the cause of this discordance is biological or just a failure to infer the correct tree.

First, and unsurprisingly, we confirmed that shorter internal branches tend to be associated with higher levels of gene tree discordance because of tree inference error (Fig. 4a). Essentially, the shorter the time window during which a population can accumulate substitutions before a speciation event takes place, the weaker the phylogenetic signal will be, and the harder it will be to recover the correct relationship among the descendant taxa. Second, we chose one gene tree to be the species tree (as done in Goldstein et al. 2015) at random, and computed the amount of convergence in the remaining sequences. We were again able to recover a pattern of diminishing convergence with increasing evolutionary distances (Fig. 4b). As mentioned above, this study also found that only parallel substitutions showed a decline with phylogenetic divergence, and not convergent substitutions. Therefore, even when biological phenomena such as ILS or introgression do not occur, the weak phylogenetic signal characteristic of short internal branches can still lead to discordant gene trees because of errors in phylogenetic reconstruction, and can translate to higher convergence among more closely related species.

**Figure 4.**
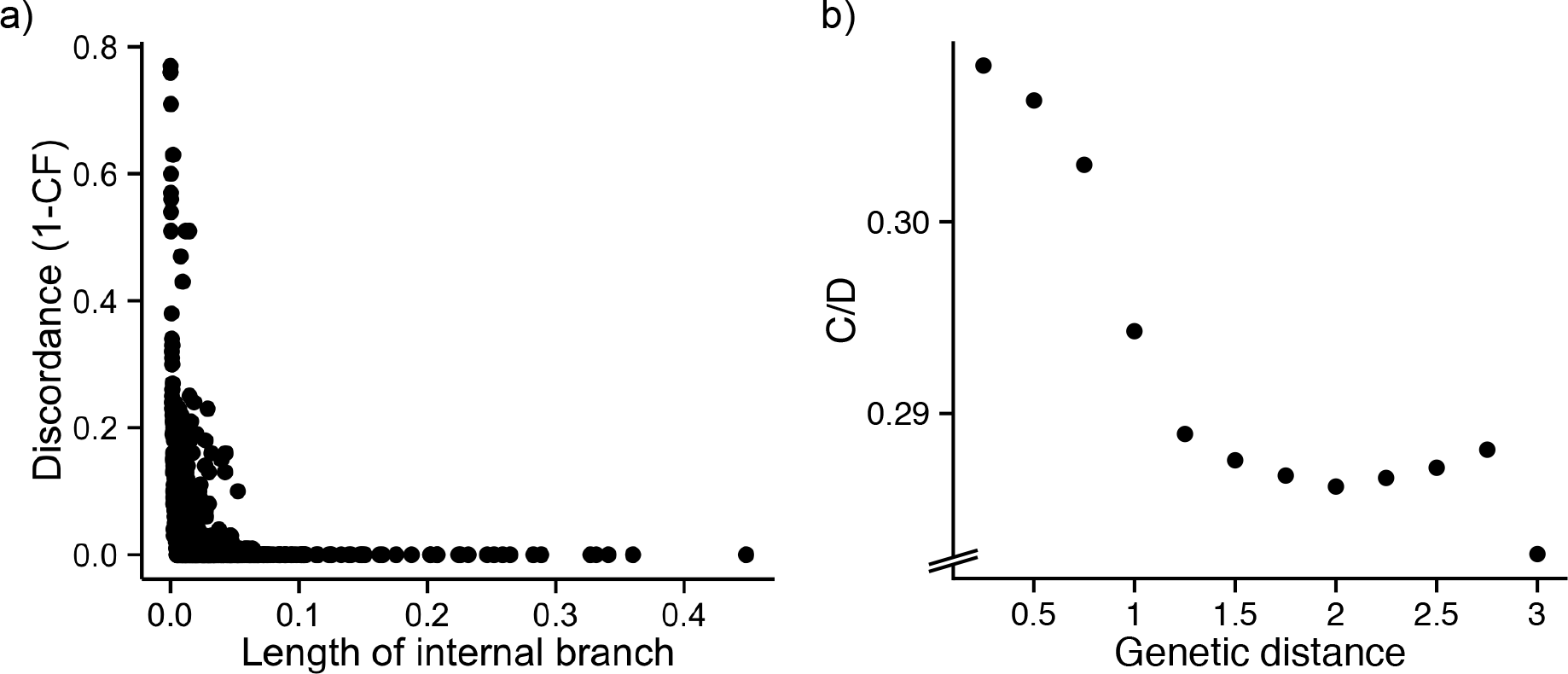
Effect of error in species tree inference on estimates of convergence. a) Discordance of nodes as a function of the length of the subtending, branch., based on data simulated along the mammalian tree from Goldstein et al. (2015). Discordance is measured as 1 – the concordance factor (CF) of the node. b) Convergence as a function of genetic distance for pairs of branches in the data simulated in part a).

## Discussion

The search for the molecular basis of convergence is currently surrounded by enthusiasm, as evidenced by the recent surge in empirical case studies within and across clades (e.g., Feldman et al. 2012; Parker et al. 2013; Projecto-Garcia et al. 2013; Goldstein et al. 2015; Zou and Zhang 2015a). These empirical studies have also stimulated a number of simulation and theoretical investigations (e.g., Pollock et al. 2012; Shah et al. 2015; McCandlish et al. 2016), as well as literature reviews (e.g., Stern 2013; Rosenblum et al. 2014; Storz 2016). This interest stems from the fact that understanding the molecular causes of convergence can provide a link between genotype and adaptive phenotypes, and more generally can reveal how repeatable and predictable adaptive evolution can be.

Large-scale inferences about the repeatability and predictability of evolution at the molecular level are generally made by counting convergent substitutions across a phylogeny (e.g., Bazykin et al. 2007; Naumenko et al. 2012; Goldstein et al. 2015; Zou and Zhang 2015a). These substitutions are identified by performing ancestral state reconstruction on a species tree, enabling researchers to polarize substitutions along specific branches. Studies employing this procedure have uncovered similar patterns: more closely related species exhibit higher levels of convergence than more distant ones. But what drives the decay of convergence with time?

Positive selection is perhaps the most investigated candidate driver of molecular convergence levels across clades, as it is expected to make convergent evolution significantly more likely (Orr 2005). Nonetheless, recent studies seem to agree that positive selection is not needed to explain the excess convergence observed among closely related species (Rokas and Carroll 2008; Goldstein et al. 2015; Zou and Zhang 2015a). Instead, convergent substitutions are thought to occur more frequently shortly after species divergence because of epistasis. This model proposes that the set of epistatically interacting sites is expected to be more similar in closely related species, thus imposing similar constraints on these sites’ otherwise independent evolution. These shared constraints are what make convergence more likely. As species diverge, epistatic interactions also diverge, which then makes convergence less likely to occur (Goldstein et al. 2015). This logic has been incorporated in models of protein evolution that allow substitution rates to vary across sites and time (Pollock et al. 2012) and whose predictions fit empirical data (Goldstein et al. 2015)

In the present study, we further extend the list of possible causes for the decay of convergence over time by demonstrating that it is also possible without epistasis. We show that in the presence of gene tree discordance, performing ancestral state reconstruction and mapping substitutions on a single species tree leads to the phenomenon of SPILS (Mendes and Hahn 2016), spuriously inflating counts of parallel substitutions at shorter genetic distances. We provide three lines of evidence that the pattern of diminishing convergent substitutions observed in real data is at least in part caused by SPILS. First, we show that when ILS is controlled for by conditioning on distinct ancestral states, convergence levels do not change with time. Second, we demonstrate that synonymous substitutions (in which epistasis should not be prevalent) also display the same pattern of diminishing convergence with greater phylogenetic distance. Finally, we uncover the same pattern of diminishing convergence over as a function of time in sequences simulated under the simple assumptions of the coalescent model, where ILS is present but epistasis is not.

Our interpretation of previous results is in accord with a growing appreciation for the ubiquity of discordance in phylogenomic datasets (Mallet et al. 2016). The discordance in eukaryotic genomes is largely due to incomplete lineage sorting and introgression, with both processes likely acting in most clades (e.g., Cui et al. 2013; Martin et al. 2013; Fontaine et al. 2015; Pease et al. 2016). Although early studies of genome-wide convergence recognized that discordance could be a problem (Rokas and Carroll 2008), the magnitude of the contribution of ancestral polymorphism and hybridization to incongruence among genes was not yet appreciated. More recent work has placed this discordance at the forefront of phylogenetic research, with a proliferation of new methods for inferring a single species tree in the face of discordance (e.g., Drummond and Rambaut 2007; Liu 2008; Larget et al. 2010; Liu et al. 2010; Bryant et al. 2012; Mirarab and Warnow 2015; Yang 2015). However, methods that can deal with discordance when analyzing patterns of molecular evolution have not kept pace, as most ancestral state reconstruction methods are still applied to only a single species tree, often with dire consequences for downstream inferences (Hahn and Nakhleh 2016).

We hope that the results presented here further highlight the need for discordance-aware phylogenetic methods. While some such methods exist (e.g., Vernot et al. 2008; Rasmussen and Kellis 2012; Maio et al. 2013; Maio et al. 2015), they are not in wide use, and hemiplasy appears to affect studies of nucleotide substitution rates and positive selection (Mendes and Hahn 2016), convergence (this study), and gene duplication and loss (Hahn 2007). Because single substitutions on discordant trees can be resolved on the species tree as either convergence or reversion (Mendes and Hahn 2016), it is likely that patterns of diminishing reversals over time may also be affected by SPILS (Naumenko et al. 2012; McCandlish et al. 2016). Due to the widespread occurrence of discordance across the tree of life – in all genomic compartments – it is likely that hemiplasy has affected many other genomic features as well. A complete reckoning of these effects is likely to encompass all aspects of molecular evolution.

## Materials and Methods

### Quantifying molecular convergence in primate coding sequences

Coding sequence alignments of orthologs from 12 primate species (human, chimpanzee, gorilla, orangutan, gibbon, rhesus macaque, cynomolgus macaque, baboon, green monkey, marmoset, squirrel monkey, and bushbaby) and an outgroup (tree shrew) were downloaded using the UCSC Table Browser tool (Karolchik et al. 2004). After removal of gaps and ambiguous codons, alignments were kept if their lengths were a multiple of 3, canonical start and stop codons were present in the human sequence, the alignment had at least 500 codons after removal of start and stop codons, and if they were the longest isoform when multiple isoforms existed for a gene. The remaining 5,264 alignments (totaling 15,198,099 sites) were used in downstream analyses.

A species tree for these 13 species was inferred (i) from a concatenated alignment using PhyML (Guindon and Gascuel 2003), (ii) with majority consensus using the PHYLIP package (Felsenstein 1989), and (iii) with MP-EST, a coalescent-based method (Liu et al. 2010). All inferred species trees had the same topology. We used this topology and branch lengths from the species tree obtained through concatenation to reconstruct ancestral states and to calculate pairwise genetic distances, both of which were needed to measure molecular convergence.

For each alignment we measured molecular convergence in a pairwise fashion by counting how many times substitutions along two non-sister branches that did not share the same phylogenetic path (i.e., we excluded comparisons involving branches that shared the same ancestral node, or two branches along the same lineage) resulted in the same nucleotide state in both descendant nodes. A substitution was mapped to a branch if at a particular site the nucleotide state at the ancestral node differed from that of the descendant node. Mapping of substitutions thus required ancestral state reconstruction, which we performed on all alignments and along the same species tree topology using the program codeml or baseml from the PAML suite (Yang 2007).

We further classified molecular convergence as parallel if the ancestral states of both ancestral nodes were the same, and as convergent if not. We normalized these two measures of molecular convergence – counts of parallel and convergent sites – by dividing them by the correspondent counts of divergent sites, respectively (in divergent sites, the focal pair of species have different nucleotide states). These two normalized measures of convergence were both referred to as C/D (counts of convergent and/or parallel substitutions normalized by counts of divergent substitutions; Goldstein et al. 2015).

In order to control for epistasis, we also analyzed the primate coding data subset containing just synonymous substitutions. Convergent and parallel changes were counted for synonymous substitutions following the same procedure described above.

Finally, we calculated the genetic distance between two taxa by adding up the lengths of all branches along the phylogenetic path connecting them. We used branch lengths from the species tree obtained from the concatenated alignment. Because at shorter genetic distances C/D had a much larger variance (as a result of C/D being a ratio of counts), we binned the data to obtain larger counts from pairs of less diverged species. The unbinned data are shown in Supplementary Figure 1.

### Quantifying molecular convergence in simulated sequences from the primate tree

We employed the primate species tree obtained with MP-EST (Liu et al. 2010) to conduct 5,000 coalescent simulations of 1 kb-long DNA sequences (*N*_e_=10,000, μ= 10_e-_7 substitutions per site) using eggcoal, a simulator from the egglib package (Mita and Siol 2012). MP-EST returns a species tree in coalescent units, which is why it was used for input into eggcoal. Internal branch lengths from the MP-EST tree were used as the initial values for simulations, but we adjusted these values slightly to produce approximately the same amount of concordance at each node (measured with concordance factors) observed in the primate tree. Divergence times of sister terminal branches were set to the average length of both taxa in the primate tree; the remaining divergence times were subsequently defined by adding the corresponding adjusted internal branch lengths. Quantification of molecular convergence and genetic distances was then performed as described above for primate coding sequences.

### Quantifying tree inference error and molecular convergence in simulated mitochondrial data

We employed the 629-species phylogeny from Goldstein et al. (2015) to simulate 100 1-kb-long coding sequences under the JC69 model (Jukes and Cantor 1969) with evolver, a program from the PAML suite (Yang 2007). Gene trees were inferred under the same model with PhyML (Guindon and Gascuel 2003) and their topologies were used to calculate the amount of discordance (as the reciprocal of the concordance factor) at each node of the phylogeny. By arbitrarily assuming one of the gene trees to be the species tree, we calculated the amount of molecular convergence observed in the remaining 99 alignments as described above.

## ACKNOWLEDGEMENTS

We thank the members of the Hahn lab for helpful discussions. This work was supported by National Science Foundation grant DEB-1136707.

## Supplementary Figure Captions

**Figure S1.**
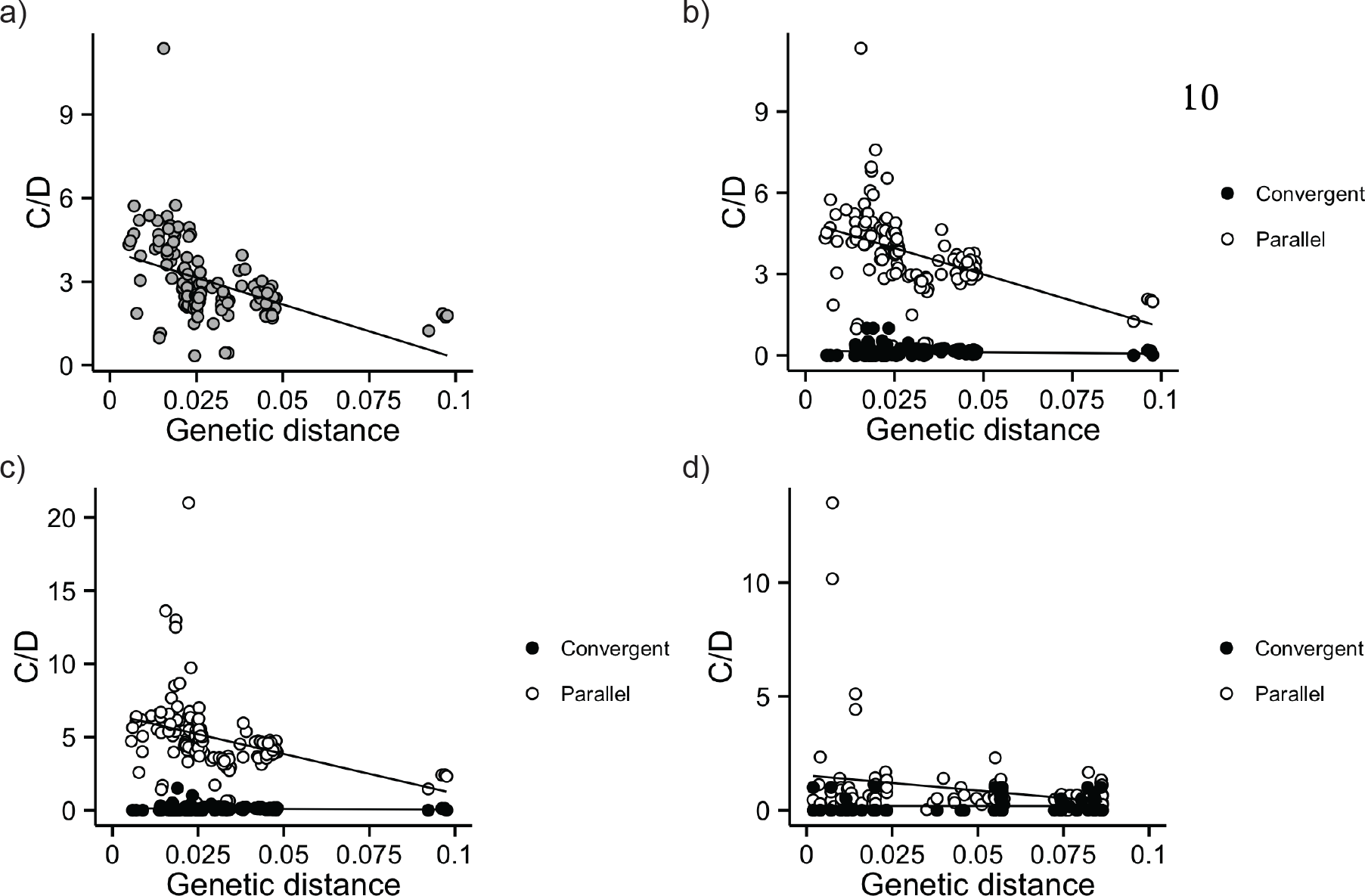
Convergence as a function of genetic distance for all pairs of species (unbinned data). a) All substitutions. b) All parallel and convergent substitutions. c) Synonymous parallel and convergent substitutions only. d) Sequences simulated on the primate species tree under the multi-species coalescent.

## REFERENCES

Avise, JC., Robinson TJ. 2008. Hemiplasy: a new term in the lexicon of phylogenetics. Systematic Biology 57:503–507.

Bazykin, GA., Kondrashov, FA., Brudno, M., Poliakov, A., Dubchak, I., Kondrashov AS. 2007. Extensive parallelism in protein evolution. Biology Direct 2:20.

Brawand, D., Wagner, CE., Li, YI., Malinsky, M., Keller, I., Fan, S., Simakov, O., Ng, AY., Lim, ZW., Bezault, E., et al. 2014. The genomic substrate for adaptive radiation in African cichlid fish. Nature 513:375–381.

Bryant, D., Bouckaert, R., Felsenstein, J., Rosenberg, N., RoyChoudhury A. 2012. Inferring species trees directly from biallelic genetic markers: bypassing gene trees in a full coalescent analysis. Molecular Biology and Evolution 29:1917–1932.

Christin, P., Salamin, N., Muasya, A., Roalson, E., Russier, F., Besnard G. 2008. Evolutionary switch and genetic convergence on rbcL following the evolution of C4 photosynthesis. Molecular Biology and Evolution 25:2361–2368.

Coyne, JA., Orr HA. 1989. Patterns of speciation in Drosophila. Evolution 43:362–381.

Cui, R., Schumer, M., Kruesi, K., Walter, R., Andolfatto, P., Rosenthal GG. 2013. Phylogenomics reveals extensive reticulate evolution in Xiphophorus fishes. Evolution 67:2166–2179.

Dobler, S., Dalla, S., Wagschal, V., Agrawal AA. 2012. Community-wide convergent evolution in insect adaptation to toxic cardenolides by substitutions in the, Na.,K-ATPase. Proceedings of the National Academy of Sciences of the United States of America 109:13040–13045.

Drummond, AJ., Rambaut A. 2007. BEAST: Bayesian evolutionary analysis by sampling trees. BMC Evolutionary Biology 7:214.

Feldman, CR., Brodie, Jr. ED., Pfrender ME. 2012. Constraint shapes convergence in tetrodotoxin-resistant sodium channels of snakes. Proceedings of the National Academy of Sciences of the United States of America 109:4556–4561.

Felsenstein J. 1989. PHYLIP – Phylogeny Inference Package (version 3.2). Cladistics 5:164–166.

Fontaine, M., Pease, JB., Steele, A., Waterhouse, R., Neafsey, D., Sharakhov, I., Jiang, X., Hall, A., Catteruccia, F., Kakani, E., et al. 2015. Extensive introgression in a malaria vector species complex revealed by phylogenomics. Science 347:1258524.

Foote, AD., Liu, Y., Thomas, GW., Vinar, T., Alfoldi, J., Deng, J., Dugan, S., van Elk, CE., Hunter, ME., Joshi, V., et al. 2015. Convergent evolution of the genomes of marine mammals. Nature Genetics 47:272–275.

Goldstein, RA., Pollard, ST., Shah, SD., Pollock DD. 2015. Nonadaptive smino acid convergence rates decrease over time. Molecular Biology and Evolution 32:1373–1381.

Guindon, S., Gascuel O. 2003. A, simple, fast, and accurate algorithm to estimate large phylogenies by maximum likelihood. Systematic Biology 52:696–704.

Hahn, MW., Nakhleh L. 2016. Irrational exuberance for resolved species trees. Evolution 70:7–17.

Hahn MW. 2007. Bias in phylogenetic tree reconciliation methods: implications for vertebrate genome evolution. Genome Biology 8:R141.

Jarvis, ED., Mirarab, S., Aberer, AJ., Li, B., Houde, P., Li, C., Ho, SY., Faircloth, BC., Nabholz, B., Howard, JT., et al. 2014. Whole-genome analyses resolve early branches in the tree of life of modern birds. Science 346:1320–1331.

Jukes, TH., Cantor CR. 1969. Evolution of protein molecules. In: New York: Academic Press. p. 21–132.

Karolchik, D., Hinrichs, AS., Furey, TS., Roskin, KM., Sugnet, CW., Haussler, D., Kent JW. 2004. The UCSC table browser data retrieval tool. Nucleic Acids Research 32:D493–496.

Larget, B., Kotha, S., Dewey, C., Ane C. 2010. BUCKy: Gene tree/species tree reconciliation with Bayesian concordance analysis. Bioinformatics 26:2910–2911.

Liu, L., Yu, L., Edwards SV. 2010. A maximum pseudo-likelihood approach for estimating species trees under the coalescent model. BMC Evolutionary Biology 10:302.

Liu L. 2008. BEST: Bayesian estimation of species trees under the coalescent model. Bioinformatics 24:2542–2543.

Maddison WP. 1997. Gene trees in species trees. Systematic Biology46:523–536.

Maio, N., Schlotterer, C., Kosiol C. 2013. Linking great apes genome evolution across time scales using polymorphism-aware phylogenetic models. Molecular Biology and Evolution 30:2249–2262.

Maio, N., Schrempf, D., Kosiol C. 2015. PoMo: an allele frequency-based approach for species tree estimation. Systematic Biology 64:1018–1031.

Mallet, J., Besansky, N., Hahn MW. 2016. How reticulated are species? Bioessays 38:140–149.

Manceau, M., Domingues, VS., Linnen, CR., Rosenblum, E., Hoekstra HE. 2010. Convergence in pigmentation at multiple levels: mutations, genes and function. Philosophical Transactions of the Royal Society B 365:2439–2450.

Martin, SH., Dasmahapatra, KK., Nadeau, NJ., Salazar, C., Walters, JR., Simpson, F., Blaxter, M., Manica, A., Mallet, J., Jiggins CD. 2013. Genome-wide evidence for speciation with gene flow in Heliconius butterflies. Genome Research 23:1817–1828.

McCandlish, DM., Shah, P., Plotkin JB. 2016. Epistasis and the dynamics of reversion in molecular evolution. Genetics. DOI: 10.1534/genetics.116.188961.

Mendes, FK., Hahn MW. 2016. Gene tree discordance causes apparent substitution rate variation. Systematic Biology. DOI::10.1093/sysbio/syw018.

Mirarab, S., Warnow T. 2015. ASTRAL-II: coalescent-based species tree estimation with many hundreds of taxa and thousands of genes. Bioinformatics 31:i44–i52.

Mita, S., Siol M. 2012. EggLib: processing, analysis and simulation tools for population genetics and genomics. BMC Genetics 13:27.

Naumenko, SA., Kondrashov, AS., Bazykin GA. 2012. Fitness conferred by replaced amino acids declines with time. Biology Letters 8:825–828.

Oliver JC. 2013. Microevolutionary processes generate phylogenomic discordance at ancient divergences. Evolution 67:1823–1830.

Orr HA. 2005. The probability of parallel evolution. Evolution 59:216–220.

Pamilo, P., Nei M. 1988. Relationships between gene trees and species trees. Molecular Biology and Evolution 5:568–583.

Parker, J., Tsagkogeorga, G., Cotton, JA., Liu, Y., Provero, P., Stupka, E., Rossiter SJ. 2013. Genome-wide signatures of convergent evolution in echolocating mammals. Nature 502:228–231.

Pease, JB., Haak, DC., Hahn, MW., Moyle LC. 2016. Phylogenomics reveals three sources of adaptive variation during a rapid radiation. PLoS Biology 14:e1002379.

Pollard, DA., Iyer, VN., Moses, AM., Eisen MB. 2006. Widespread discordance of gene trees with species tree in Drosophila: evidence for incomplete lineage sorting. PLoS Genetics 2:1634–1647.

Pollock, DD., Thiltgen, G., Goldstein RA. 2012. Amino acid coevolution induces an evolutionary Stokes shift. Proceedings of the National Academy of Sciences of the United States of America 109:E1352–1359.

Povolotskaya, IS., Kondrashov FA. 2010. Sequence space and the ongoing expansion of the protein universe. Nature 465:922–926.

Projecto-Garcia, J., Natarajan, C., Moriyama, H., Weber, RE., Fago, A., Cheviron, ZA., Dudley, R., McGuire, JA., Witt, CC., Storz JF. 2013. Repeated elevational transitions in hemoglobin function during the evolution of Andean hummingbirds. Proceedings of the National Academy of Sciences of the United States of America 110:20669–20674.

Rasmussen, M., Kellis M. 2012. Unified modeling of gene, duplication, loss, and coalescence using a locus tree. Genome Research 22:755–765.

Rogozin, IB., Thomson, K., Csuros, M., Carmel, L., Koonin EV. 2008. Homoplasy in genome-wide analysis of rare amino acid replacements: the molecular-evolutionary basis for Vavilov’s law of homologous series. Biology Direct 3:7.

Rokas, A., Carroll SB. 2008. Frequent and widespread parallel evolution of protein sequences. Molecular Biology and Evolution 25:1943–1953.

Rosenblum, EB., Parent, CE., Brandt EE. 2014. The molecular basis of phenotypic convergence. Annual Review of, Ecology, Evolution, and Systematics 45:203–226.

Scally, A., Dutheil, JY., Hillier, LW., Jordan, GE., Goodhead, I., Herrero, J., Hobolth, A., Lappalainen, T., Mailund, T., Marques-Bonet, T., et al. 2012. Insights into hominid evolution from the gorilla genome sequence. Nature 483:169–175.

Shah, P., McCandlish, DM., Plotkin JB. 2015. Contingency and entrenchment in protein evolution under purifying selection. Proceedings of the National Academy of Sciences of the United States of America 112:E3226–3235.

Stern DL. 2013. The genetic causes of convergent evolution. Nature Reviews Genetics 14:751–764.

Storz JF. 2016. Causes of molecular convergence and parallelism in protein evolution. Nature Reviews Genetics 17:239–250.

Suh, A., Smeds, L., Ellegren H. 2015. The dynamics of incomplete lineage sorting across the ancient adaptive radiation of neoavian birds. PLoS Biology 13: e1002224.

Thomas, GW., Hahn MW. 2015. Determining the null model for detecting adaptive convergence from genomic data: a case study using echolocating mammals. Molecular Biology and Evolution 32:1232–1236.

Vernot, B., Stolzer, M., Goldman, A., Durand D. 2008. Reconciliation with non-ninary species trees. Journal of Computational Biology 15:981–1006.

Yang Z. 2007. PAML 4: Phylogenetic Analysis by Maximum Likelihood. Molecular Biology and Evolution 24:1586–1591.

Yang Z. 2015. The BPP program for species tree estimation and species delimitation. Current Zoology 61:854–865.

Zhang, G., Li, C., Li, Q., Li, B., Larkin, DM., Lee, C., Storz, JF., Antunes, A., Greenwold, MJ., Meredith, RW., et al. 2014. Comparative genomics reveals insights into avian genome evolution and adaptation. Science 346:1311–1320.

Zhang, J., Kumar S. 1997. Detection of convergent and parallel evolution at the amino acid sequence level. Molecular Biology and Evolution 14:527–536.

Zhen, Y., Aardema, ML., Medina, EM., Schumer, M., Andolfatto P. 2012. Parallel molecular evolution in an herbivore community. Science 337:1634–1637.

Zou, Z., Zhang J. 2015a. Are convergent and parallel amino acid substitutions in protein evolution more prevalent than neutral expectations? Molecular Biology and Evolution 32:2085–2096.

Zou, Z., Zhang J. 2015b. No genome-wide protein sequence convergence for echolocation. Molecular Biology and Evolution 32:1237–1241.

